# An AlphaFold guided model for the evolution of the CaMKII interactome

**DOI:** 10.1101/2025.08.21.670683

**Authors:** Salman T. Khawaja, Maham Hamid, Thomas S. Reese, Shahid Khan

**Author notes:** Deceased (Thomas S. Reese).

## Abstract

The neuronal functions of mammalian calcium calmodulin (Ca^2+^.CAM) dependent protein kinase II (CaMKII) are orchestrated by an interactome of multiple CaMKII-protein interactions when Ca^2+^.CAM opens the kinase domain to bind Ca^2+^ response regulators and substrates to its regulatory helix and C-lobe, respectively. We analyzed over forty 3D-atomic structures and models of CaMKII-target complexes to track the evolution of the neuronal CaMKII interactome over three model organisms and the early metazoan Trichoplax adhaerans. We report the conservation of the molecular interactome based on binding surface overlap, energy frustration and fold evolution. Transcriptome databases informed on Ca^2+^ CaMKII response regulation and colocalization with substrates. The activation machinery was invariant, but the co-expression of CaMKII and its substrates relative to the mammalian α isoform was progressively reduced in simpler organisms. We propose CaMKII architecture for autoinhibition, Ca^2+^ response and substrate association was formed for neurosecretion, then specialized for synaptic signalling with the α-isoform.

## Introduction

The calcium calmodulin (Ca^2+^.CAM) dependent protein kinase II (CaMKII) has a fundamental role in coupling electrical activity to phosphorylation circuits via Ca^2+^ signals ^1,2^ and remodeling of structural scaffolds ^3,4^. In mammals, it is expressed in a broad range of tissues with diverse functions. A common CaMKII subunit architecture has evolved across species (**Fig. 0a**). Gene duplications have created four mammalian CaMKII isoforms (α, β, δ, γ), each with multiple alternatively spliced linker variants. The α isoform is dedicated for neuronal function. In humans, CaMKIIα mutations result in intellectual disabilities with defects in neuronal development as well as electrophysiological circuitry ^5-7^. The clinical data have been supplemented by biochemical, live-cell imaging, and electrophysiological studies on mammalian models, mainly rodents ^8,9^. These studies have mapped the cellular and biochemical basis for the role of CaMKII in long-term potentiation, a phenomenon associated with learning and memory ^1,10,11^. Other mammalian CaMKII isoforms (δ, γ) have important roles in cardiac function ^12^ and urinary filtration ^13^.

**Figure 0:**
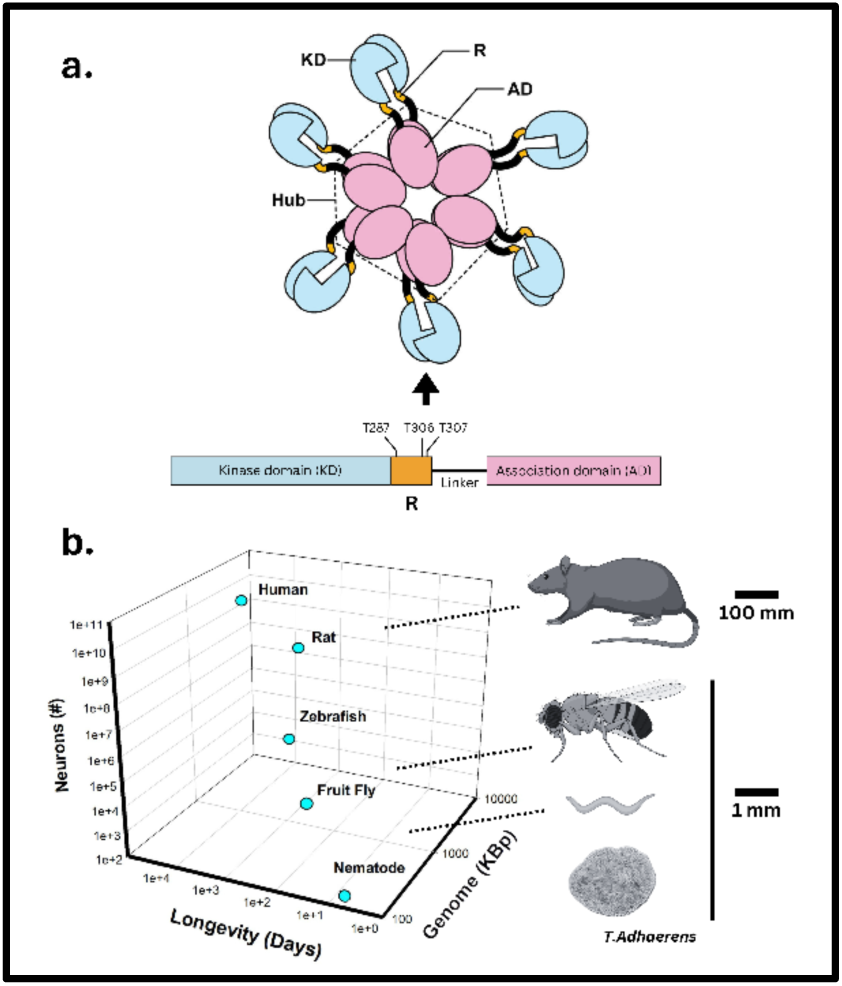
Size and complexity of organisms with CaMKII: **(a).** The CaMKII holoenzyme consists of 12-16 copies of the individual subunits arranged as a two-ring stack. The C-terminal association domains (ADs) form a two-stack ringed hub, surrounded by the peripheral N-terminal kinase domains (KDs) ^27^. Flexible linkers with variable length and sequence connect the KDs to the AD hub ^28^. The autoinhibitory R segment and its activating (αT286) and inhibitory (αT305.T306) autophosphorylation sites localise to the KD C-terminus. While polymorphs with variable stoichiometries coexist within single populations, the hub interfaces are conserved from mammals to early metazoa ^29^. **(b).** The evolution of complexity. Plot of the correlation between genome size, longevity and neuronal complexity. Rodents, the common models of memory, range in age from 9-24 months. Model invertebrates have lifetimes that range from days to weeks. Fruit flies (40-50 days); worms (3-5 days). Long-term memory is absent in these models, but there is learning and habituation memory. The placozoan, T. adhaerans, replicates by asexual fission and has no known CaMKII function.

CaMKII is an ancient enzyme present in early metazoans. The placozoan *Trichoplax Adhaerans*, an organism with neuropeptides but not neurons, is related to the cnidarian *Nematostella vectensis* ^14^, with a single CaMKII encoded for unknown functions. The insect *Drosophila melanogaster* and nematode *Caenorhabditis elegans* also have a single CaMKII gene. Their nervous systems are wired differently, but they have synapses ^15^. They have been studied as models for learning and memory (see reviews (*D. melanogaster* ^16-18^, *C. elegans* ^19-21^))

The overarching issue raised in organisms with single CaMKII genes is the nature of the functional spectrum in the absence of isoforms dedicated for specialized functions, accentuated by the absence of alternatively spliced variants for *T. Adhaerans* CaMKII. It has been problematic to relate the roles of primate CaMKIIs in complex cognitive functions to non-primate models from behavioral and physiological assays alone ^9^. The problem is compounded for invertebrates and early metazoa where shortened life-spans, small size, reduced body-plan and genome complexity make it a challenge to define behavioral terms such as “long-term potentiation” (**Fig. 0b**). In previous work, the emergence of the human CaMKIIα hub from primordial ancestors was mapped at the molecular level with residue coevolution^22^ and energetic frustration^23^, supplemented with phylogenetics^24^. Here, we extend this structural approach with AlphaFold (AF) atomic models ^25,26^ to frame the evolution of CaMKII interactions with its primary targets in the model invertebrates, and *T. Adhaerans* relative to the mammalian rodent model. Residue coevolution is incorporated as one input in the AF neural net algorithm ^26^.

CaMKII is switched on by Ca^2+^.CAM at optimal Ca^2+^ pulse frequencies ^30^ from the basal (CLOSED) to the activated (OPEN) state. In the CLOSED state, the autoinhibitory segment, R, occludes the target-binding groove (TBG) in the KD C-lobe. Ca^2+^.CAM dissociates R from the KD core to create the substrate-accessible OPEN state ^31^. R extends from the AD hub ^32^ for capture and αT286 trans-phosphorylation ^33^. R αT286 phosphorylation traps Ca^2+^.CAM ^34,35^, while Inhibitory αT305.T306 autophosphorylation blocks Ca^2+^.CAM association ^36,37^. CaMKII binds targets in the OPEN state to the freed R (R*_f_*), or the conserved C-lobe target-binding groove (TBG). We first analyzed the AF CaMKII basal CLOSED state model. We then partitioned the species CaMKII sequences into two – the (KD–R) and (R + AD) segments for AF prediction of OPEN state CaMKII / target complexes. In addition to Ca^2+^.CAM, R*_f_* binds protein phosphatases ^38,39^, α-actinin ^40^, the postsynaptic scaffold Maguk homolog, Camguk ^41^ and presynaptic Synapsin-1 ^42^. The synaptic Ca^2+^ response depends on the competition between these targets and Ca^2+^.CAM for the common R*_f_* Ca^2+^.CAM binding surface ^43^. The sequestration of CaMKIIα to the synaptic membrane is mainly orchestrated by TBG targets. There are crystal structures of CaMKII (KD-R) complexes with the NMDA receptor GluN2B (7UJR.pdb), a key target ^44^ for scaffold formation, AMPA GluA1, Tiam-1 GDP-GTP exchange factor, densin-180 ^45^), in addition to the inhibitor CaMKIINtide ^33^) and the FLY K^+^ channel ^46^) peptides. CaMKIIα also tethers to Ca^2+^ channels via the TBG ^47^. We built atomic (R + AD) and (KD – R) AF models for the following species: *Rattus norvegicus* (RAT). *Drosophila melanogaster* (FLY), *Caenorhabditis elegans* (WORM) and *Trichoplax adhaerans* (TRIX) - to track the organismal evolution of the CaMKII complexes and the competition between targets. We augmented our structural models with transcriptome databases to compare CaMKII and target tissue expression patterns since complex formation, hence physiological function, depends on in vivo concentration as well as affinity. We refer to this primary set of CaMKII complexes for the construction of progressively sophisticated circuitry as the CaMKII “interactome”. The CaMKII interactomes revealed the progressive evolution of CaMKII as a neuronal Ca^2+^ signal kinase during the transition from the chemical to the electrical “brain” ^48^.

## Results & Discussion

### A. The transition between CaMKII OPEN and CLOSED states

We first tracked the evolution of the CaMKII holoenzyme. Multiple sequence alignment (MSA) partitioned the CaMKII subunit into the conserved KD and the AD across species. The databases and software used for the MSA and all subsequent operations are listed in **Table 1**. The RAT CaMKIIα isoform was the benchmark since its evolution correlates with the evolution of memory ^24^. The match scores were consistent with previous reports ^24,49^ that the mammalian CaMKIIδ isoform KD and AD are most homologous with the corresponding domains in the FLY and WORM. The homology was extended to TRIX (**Fig. 1a**).

**Figure 1:**
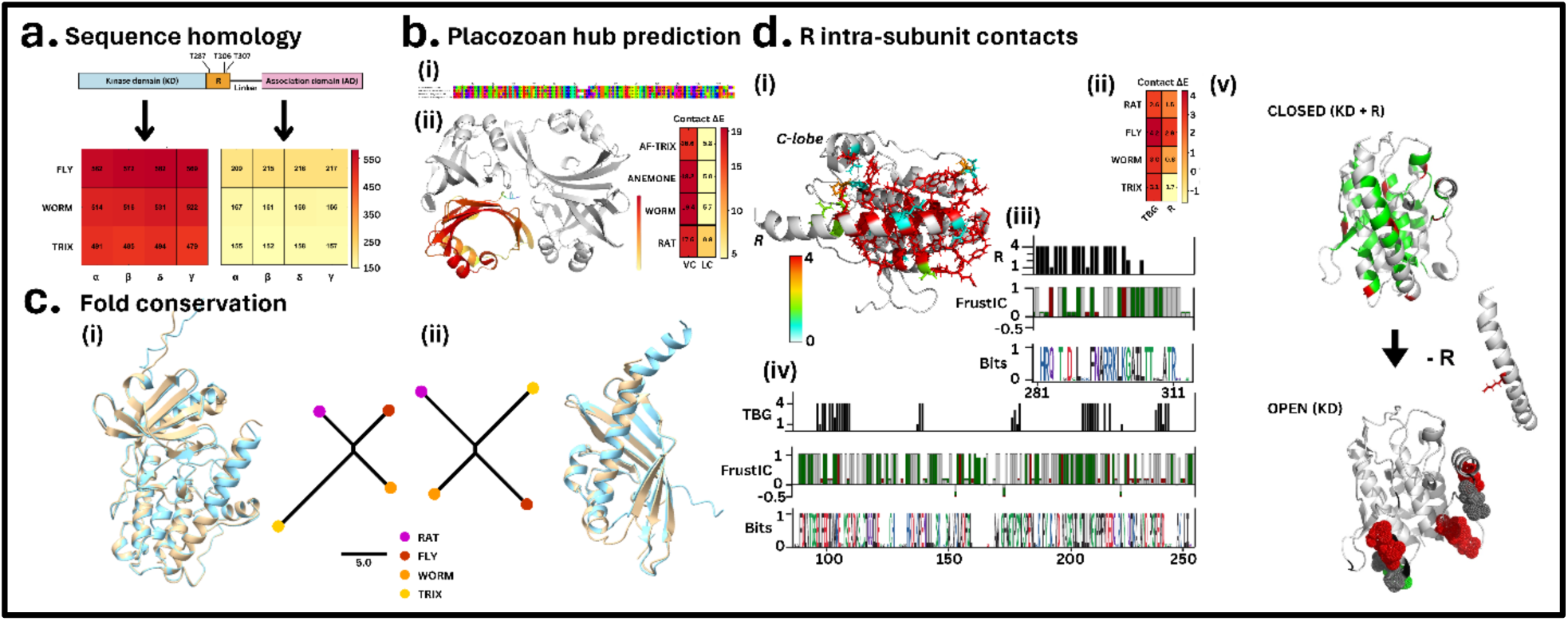
CaMKII Architecture from mammals to placozoa: **a. Sequence homology.** The sequence conservation of the invertebrate kinase (KD) and association (AD) domains relative to the four rat isoforms is shown. The match scores were a more sensitive measure than E-values of sequence variation, given the high conservation of these domains. **b**. **Placozoan hub prediction**. **(i)** Conservation of hub contact residues. **(ii)** AlphaFold (AF) model of the T. Adhaerans AD tetramer. One AD is color-coded with the pLDDT score (Bar. Low (yellow) -> High (red)). The ePISA estimates of the T. adhaerans AF-model contacts are compared against corresponding estimates for experimentally determined hub structures ^29^. **C. Fold conservation. (i) KD.** Left. Superposed human hub (3SOA.pdb (salmon)) / TRIX AF-model (cyan). Right: DALI tree of fold similarity. **(ii) AD**. Left. Superposed human hub (3SOA.pdb (salmon)) / TRIX AF-model (cyan). Right: DALI tree of fold similarity. **D. R. Intra-subunit contacts**. **(i)** 3D RAT KD extracted from the subunit model. Colors indicate R_C-lobe contact overlap between the four organisms (Bar. Low (white) -> High (red). **(ii) R-TBG Contact energetics.** The ePISA estimates for the R-TBG contact in the AF models. **(iii, iv)** Residue overlap (top), FrustratoEvo (middle) and HMM (bottom) logos for (iii) R, (iv) TBG. Frustrato Logo shows minimally frustrated (green). Highly frustrated (red) and neutral (grey) residue positions. Residues are colored by type in the HMM logo. **(v)** Energy frustration in the CLOSED versus OPEN states. In the OPEN states, only residue positions with changed assignment between the three frustration states are shown.

**Table 1:**
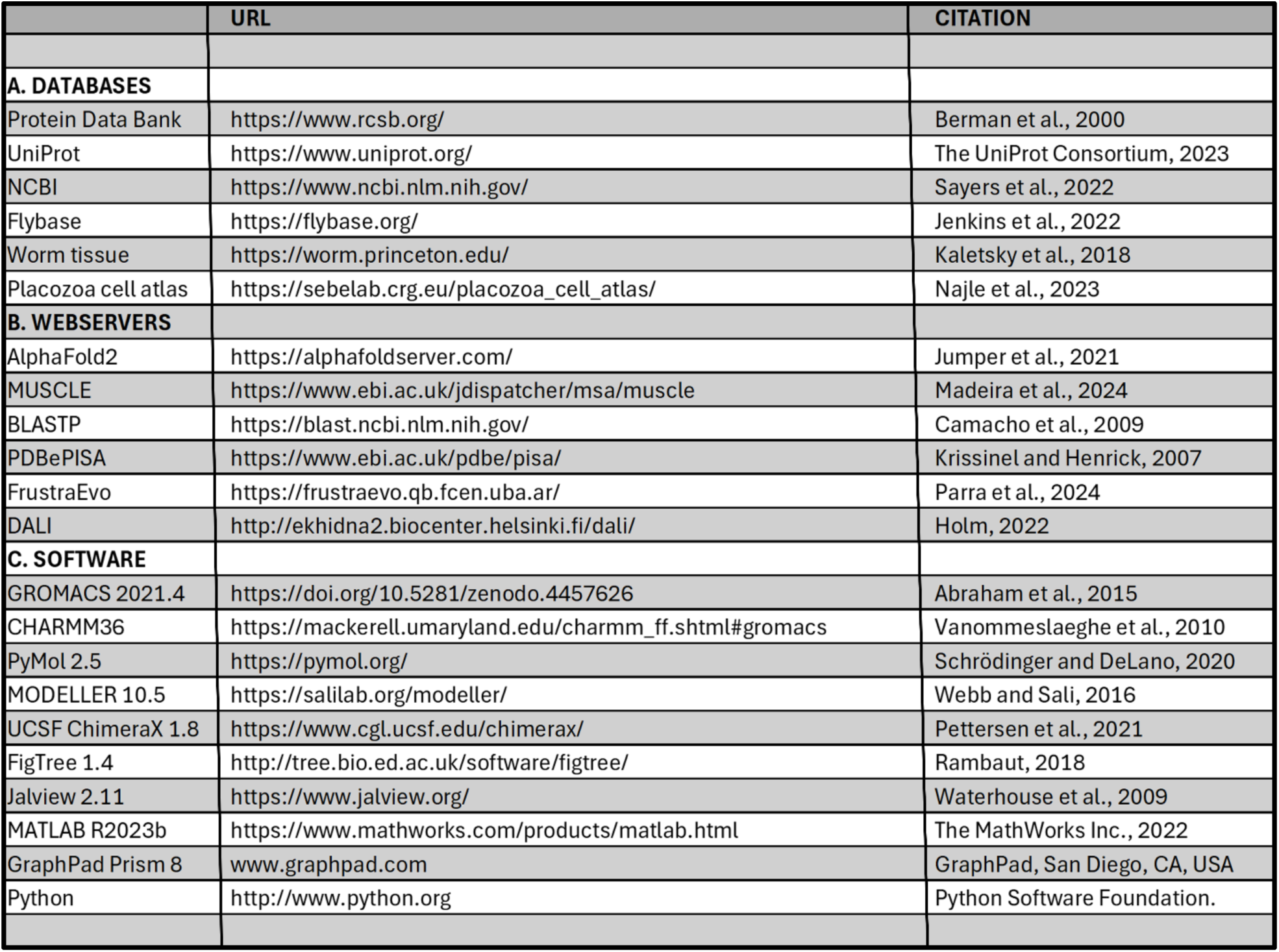
Databases, webservers and software.

|  | URL | CITATION |
| --- | --- | --- |
| **A. DATABASES** |  |  |
| Protein Data Bank | https://www.rcsb.org/ | Berman et al., 2000 |
| UniProt | https://www.uniprot.org/ | The UniProt Consortium, 2023 |
| NCBI | https://www.ncbi.nlm.nih.gov/ | Sayers et al., 2022 |
| Flybase | https://flybase.org/ | Jenkins et al., 2022 |
| Worm tissue | https://worm.princeton.edu/ | Kaletsky et al., 2018 |
| Placozoa cell atlas | https://sebelab.crg.eu/placozoa_cell_atlas/ | Najle et al., 2023 |
| **B. WEBSERVERS** |  |  |
| AlphaFold2 | https://alphafoldserver.com/ | Jumper et al., 2021 |
| MUSCLE | https://www.ebi.ac.uk/jdispatcher/msa/muscle | Madeira et al., 2024 |
| BLASTP | https://blast.ncbi.nlm.nih.gov/ | Camacho et al., 2009 |
| PDBePISA | https://www.ebi.ac.uk/pdbe/pisa/ | Krissinel and Henrick, 2007 |
| FrustraEvo | https://frustraevo.qb.fcen.uba.ar/ | Parra et al., 2024 |
| DALI | http://ekhidna2.biocenter.helsinki.fi/dali/ | Holm, 2022 |
| **C. SOFTWARE** |  |  |
| GROMACS 2021.4 | https://doi.org/10.5281/zenodo.4457626 | Abraham et al., 2015 |
| CHARMM36 | https://mackerell.umaryland.edu/charmm_ff.shtml#gromacs | Vanommeslaeghe et al., 2010 |
| PyMol 2.5 | https://pymol.org/ | Schrödinger and DeLano, 2020 |
| MODELLER 10.5 | https://salilab.org/modeller/ | Webb and Sali, 2016 |
| UCSF ChimeraX 1.8 | https://www.cgl.ucsf.edu/chimerax/ | Pettersen et al., 2021 |
| FigTree 1.4 | http://tree.bio.ed.ac.uk/software/figtree/ | Rambaut, 2018 |
| Jalview 2.11 | https://www.jalview.org/ | Waterhouse et al., 2009 |
| MATLAB R2023b | https://www.mathworks.com/products/matlab.html | The MathWorks Inc., 2022 |
| GraphPad Prism 8 | www.graphpad.com | GraphPad, San Diego, CA, USA |
| Python | http://www.python.org | Python Software Foundation. |

We next assessed the prediction accuracy of AF models for protein-protein contacts and domain fold by comparison against known crystal and cryo-EM structures (**Table 2)**. The mammalian hubs are completely described by tetramer sub-assemblies. The MSA of the hub domains for all four species suggests the inter-domain contacts are conserved, but crystal structures from *N. vectensis,* early metazoan ^14^ and algal ^50^ hubs have revealed deviations from the closed ring topology of mammalian hubs. We constructed multimer AF-models of the RAT and TRIX AD tetramer to explore this issue. The top models of the RAT CaMKIIα and the TRIX hub tetramers aligned exactly (**Fig. 1b**). Furthermore, all 25 Trix AF models did not deviate from the closed ring mammalian hub topology. The energies for vertical (VC) and lateral (LC) hub contacts were comparable across organisms. AF monomer models of the AD and KD were constructed for all four organisms (RAT, FLY, WORM, TRIX). DALI revealed fold variation between organisms for both the KD and AD models. The variations were small - superposed AF models of the RAT and TRIX aligned with RMSDs of 0.57 angstrom (KD) and 0.72 angstrom (AD), respectively (**Fig. 1c**), and did not track the phylogenetic tree, in contrast to the sequences. The TRIX CaMKII models provide the first 3D structural insights into the holoenzyme and its assemblies.

**Table 2.**
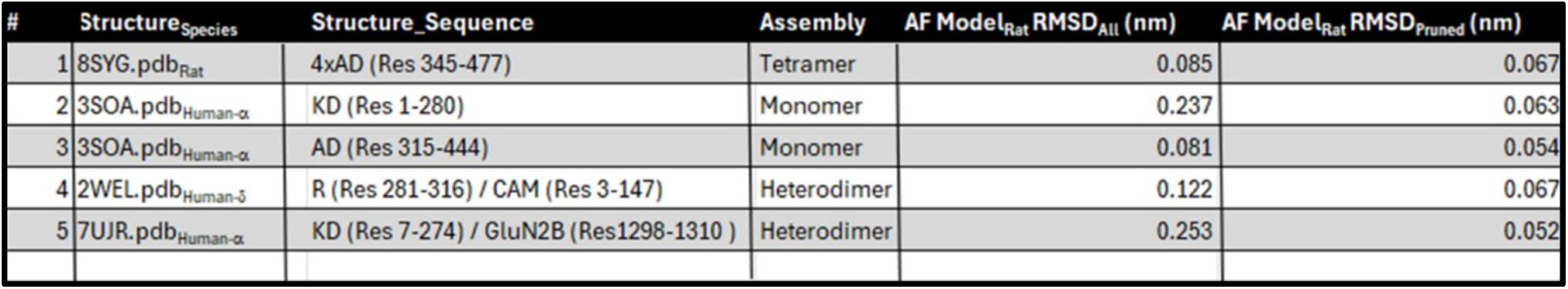
Model validation against experimental structures. The ChimeraX matchmaker tool was used to align the RAT AF models with the experimental cryo-EM (rat (8SYG.pdb)) and crystal (human (3SOA.pdb, 2WEL.pdb, 7UJR.pdb) CaMKII structures.

CaMKII targets bind predominantly to its KD. The CaMKII KD exists in at least two states, primed by Ca^2+^.CAM - the CLOSED state in which substrate access to the TBG is blocked by R docked to the C-lobe, and the OPEN state with the TBG exposed with R free (R*_f_*). The predicted AF models for the CaMKII subunit in the four organisms reflected the CLOSED human holoenzyme crystal structure (3SOA.pdb^27^) with R bound to the TBG. Comparison of the R-TBG contact energies ranked the FLY contact as the most, and the TRIX contact as the least, favorable (**Fig.1di. ii**).

The R-TBG residue contacts were conserved between organisms (**Figs. 1d.iii, iv**). They localized to R (R-1, R-2), and C-lobe segments as reported in 3SOA.pdb ^27^. We focused our study on these contacts. The energy frustration profile for the CLOSED subunit KD showed a stable core that included the bound R. The OPEN TBG FrustratoEvo organism residue profile for the two major TBG segments (TBG-1, TBG-2), when matched against the extensive HMM KD conservation logos compiled by ^24^ documented that energy frustration was not correlated with sequence conservation (**Fig. S1a**).

We next mapped the local frustration of these contacts between the top subunit (CLOSED) versus the OPEN (KD-R) AF models (**Fig.1d. v**). The energy frustration profile for the CLOSED subunit KD showed a stable core that included the bound R. The comparison of the changes in the energy frustration during the CLOSED to OPEN transition showed that the frustration of the C-lobe contact residues increased as the stabilization provided by R was removed. Similarly, the local frustration of the conserved R residue position I_293_ (RAT numbering) increased in R*_f_*.

R contacted the AD in the AF-models even though the native linker sequences were longer than the engineered short linker in the 3SOA.pdb crystal structure. The predicted R-AD contacts were heterogeneous between organisms, possibly due to the linkers. We supplemented the AF monomer prediction with multimer models with the KD and R input as separate sequences to remove possible linker-mediated effects. The multimer models predicted fewer R contacts that mostly localized to the central AB β-sheet involved in hub assembly. The multimer predictions imply the model for mammalian hub fragmentation by R peptides ^51^ may extend to lower organisms but will need experimental validation. While alternative linker splicing affects CaMKII frequency decoding and localization to cellular compartments, the disordered linkers challenge structure prediction. We note the increase in linker sequence diversity with organism complexity (**Fig. S1b**), with the homology for linker exons evaluated by decoy distributions (**Fig. S1c**), but do not study linker structure or the proposed association of cytoskeletal F-actin with a β linker exon based on peptide assays ^52^ due to the difficulty in linker structure prediction.

Our principal conclusions from the CaMKII AF-models are as follows. First, the CaMKII AD and KD sequences from placozoa to insects are most homologous to CaMKIIδ. Second, the predicted intra-domain hub and R-TBG contacts are conserved between organisms. The predicted R-AD contacts are both heterogeneous and distinct from those in the human holoenzyme structure ^27,53^. Third, local energy frustration analysis of OPEN and CLOSED states reveals that stabilization of the R and the C-lobe TBG is a common feature of CaMKII, as shown for many other complexes ^54,55^, over an extended phylogenetic range.

### B. Construction of the CaMKII interactomes

We constructed the RAT CaMKII interactome from the selected targets listed in the **Introduction**. The selection is instrumental for mammalian synaptic memory ^1,11^. We then constructed the FLY, WORM and TRIX CaMKII interactomes as detailed in **Methods** - **Target selection**. We built over 40 AF-models for the CaMKII target complexes – for validation against experimentally determined crystal structures as well as *de novo* prediction of novel complexes.

Ca^2+^.CAM binds to OPEN the CLOSED KD. The AF models of the RAT CaMKIIα subunit with and without bound Ca^2+^.CAM were compared to show that the R helix must detach from the TBG to bind Ca^2+^.CAM (**Fig. S2a**). The R / Ca^2+^.CAM contacts were indistinguishable from those reported for the crystal structure of Ca^2+^.CAM bound to human CaMKIIδ (2WEL.pdb). Therefore, AF models the Ca^2+^.CAM, and not the apo-CAM structure from the input CAM sequence.

The OPEN (R+AD) or (KD-R) CaMKII sequences were used for the AF models of bound R and TBG target complexes, respectively. Comparison of the RAT CaMKIIα AF models of the CLOSED KD in the intact subunit with the OPEN KD in the bound GluN2B / (KD-R) complex showed the rotation of helix αD (**Fig. S2b**), as reported ^27,33^. The OPEN model complexes were partitioned into groups based on cellular localization and/or function, namely (a) ion channels, (b) membrane scaffolds. (c) intracellular signalling and (d) the actin cytoskeleton. Two metrics were extracted from the complexes - the interfacial contact energy as a diagnostic for the affinity, and local energy residue frustration as an indicator of contact evolution (see **Methods**).

### C. The competition between regulatory targets for the R*f* signal module

The contact profiles, energy, and residue conservation computed from the AF-models of the OPEN R*f* target complexes are shown in **Fig. 2a-c**. The conserved central R segment *_N_*FNRRALK*_C_* has the greatest propensity summed over the top targets. This segment, known to be important for trapping CaMKII ^34^, is dedicated for association with external targets as well as the KD C-lobe (Fig. 2a). Ca^2+^.CAM binds most strongly in all four species, TRIX Ca^2+^.CAM affinity for its CaMKII R*_f_* is half of that estimated for the other organisms. There is a progressive decrease in contact strength from Ca^2+^.CAM to α-actinin then PPI. The TRIX PPI contact energetics and binding surface, while anomalous to those predicted for other organisms, were self-consistent. Ca^2+^.CAM and α-actinin both bind with EF-hand domains. The difference between them may simply reflect that the sequence input for α-actinin was limited to one, rather than two, EF-hands. The presynaptic vesicle anchor, Synapsin-I, binds with comparable strength in the RAT, FLY and WORM. It does not have a TRIX homolog (**Fig. 2b**).

**Figure 2:**
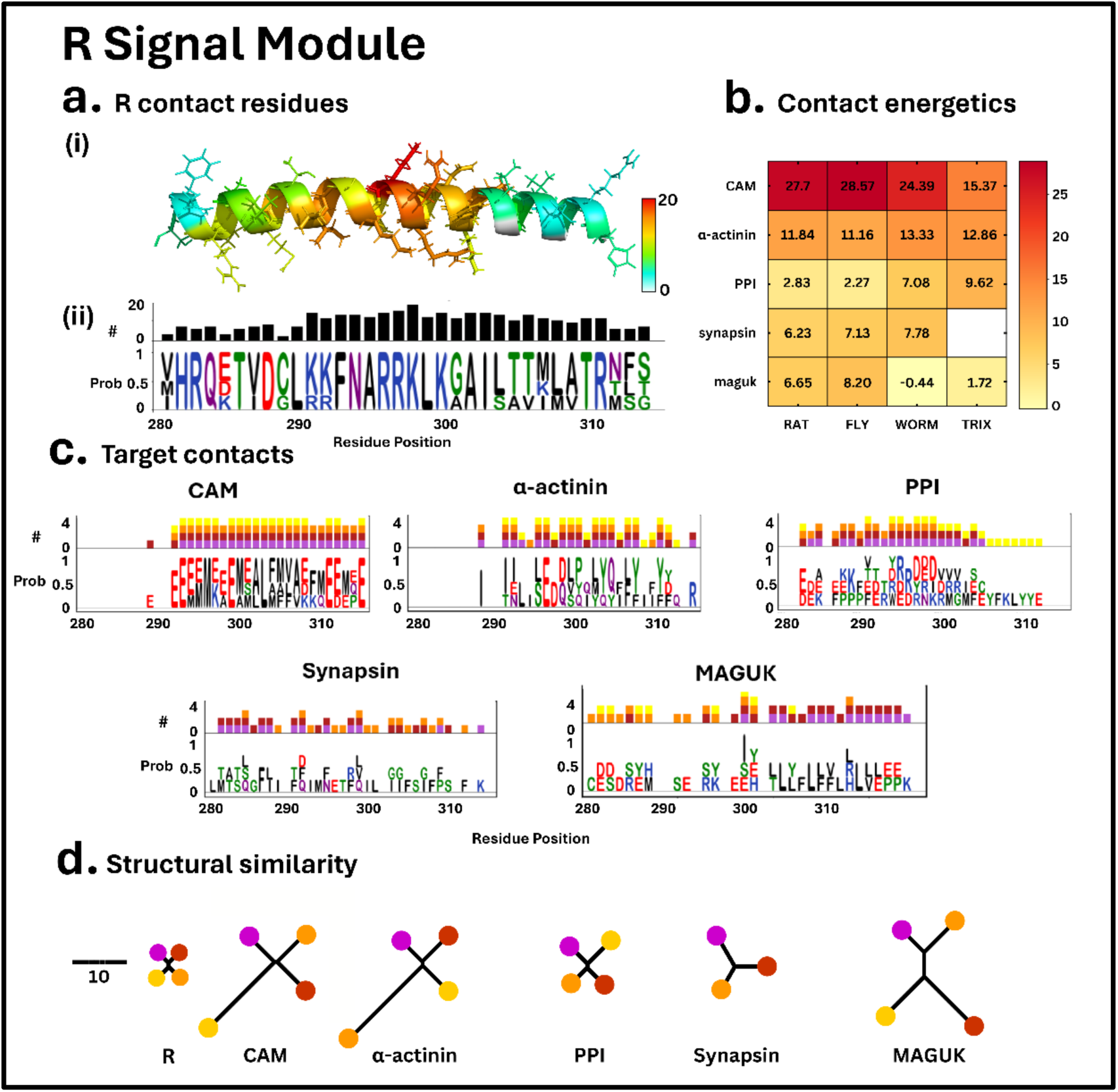
The R signal module: **a. R residue conservation and target propensity. (i)** Contact density per residue mapped onto structure**. (ii**) HMM logo. Bars (height = number of contacts per residue position) above the HMM logo indicate residue contact distribution. The maximum is 20 (4 species x 5 targets). **b. Contact energetics**. Computed by ePISA. **c. Target contact distribution** by organism. **d**. **Structural similarity.** DALI tree of the structural similarity of the contacts between organisms for the R module and its targets. Color code for organisms is as in Fig. 1c.

There is extensive overlap between the R*f* contact surface for CAM and α-actinin, as expected. The contacts extend over the entire length of R*f*. The RAT, FLY, and WORM PPI contacts localise to the N-terminal and central residues, in contrast to TRIX PPI, whose contact extends over its R*f* C-terminal as well. The extended contact results in a relatively more stable TRIX R*f* / PPI complex. The Maguk WORM and TRIX homologs make extended contacts with the R*f* C-termini, but in both cases the overall contact strength is weaker than for the corresponding RAT and FLY complexes (**Fig. 2c**).

The fold evolution of R*f* and its target contacts was measured with DALI. The R*f* helix is evolutionarily conserved relative to its targets. Among the targets, Synapsin is highly conserved between the RAT, FLY and WORM, but absent in TRIX. The PPI contact involves the central R*f* sequence segment in all four organisms, and the fold variation between the PPI domains is the least among the R*f* targets. The TRIX CaM and the WORM α-actinin folds in their R*f* complexes were outliers in their respective DALI trees. **(Fig. 2d**). The FrustratoEvo profiles for the R*f* targets are shown in (Fig. **S2c**). Contact-stabilized frustrated residue positions were observed for CAM and PPI.

### D. The TBG module target contacts have conserved overlap and comparable energetics

The mammalian TBG binds inhibitor peptides as well as substrates ^45^. The selected substrates were synaptic proteins, primarily ion channels ^44,47^. The contact overlap of these targets on the TBG groove is mapped onto the structure and plotted as a histogram (**Fig. 3a**). Contact energies were in the 4 -18 kcal/mole range (**Fig.3b**). Homologs for the mammalian CaMKIIN inhibitor peptide homologs were not present in the FLY, WORM and TRIX. The TBG contact surface was remarkably well-conserved between organisms, including TRIX (**Fig.3c**). NMDA exploits the multisubunit CaMKII assembly to form stable, higher-order scaffolds ^4,56^. It is of interest that the contact energies computed for the crystal structure (7UJR.pdb) as well as the AF-model are lower than that of the R*f* / CAM contact. This result implies that the stability of the scaffolds requires multivalent contacts of the CaMKII holoenzyme with NMDA receptors, consistent with available evidence ^57^. Alternatively, the CaMKII-NMDA complex may transition into a persistently-activated, high-affinity conformation that is not predicted by the AF algorithm.

**Figure 3:**
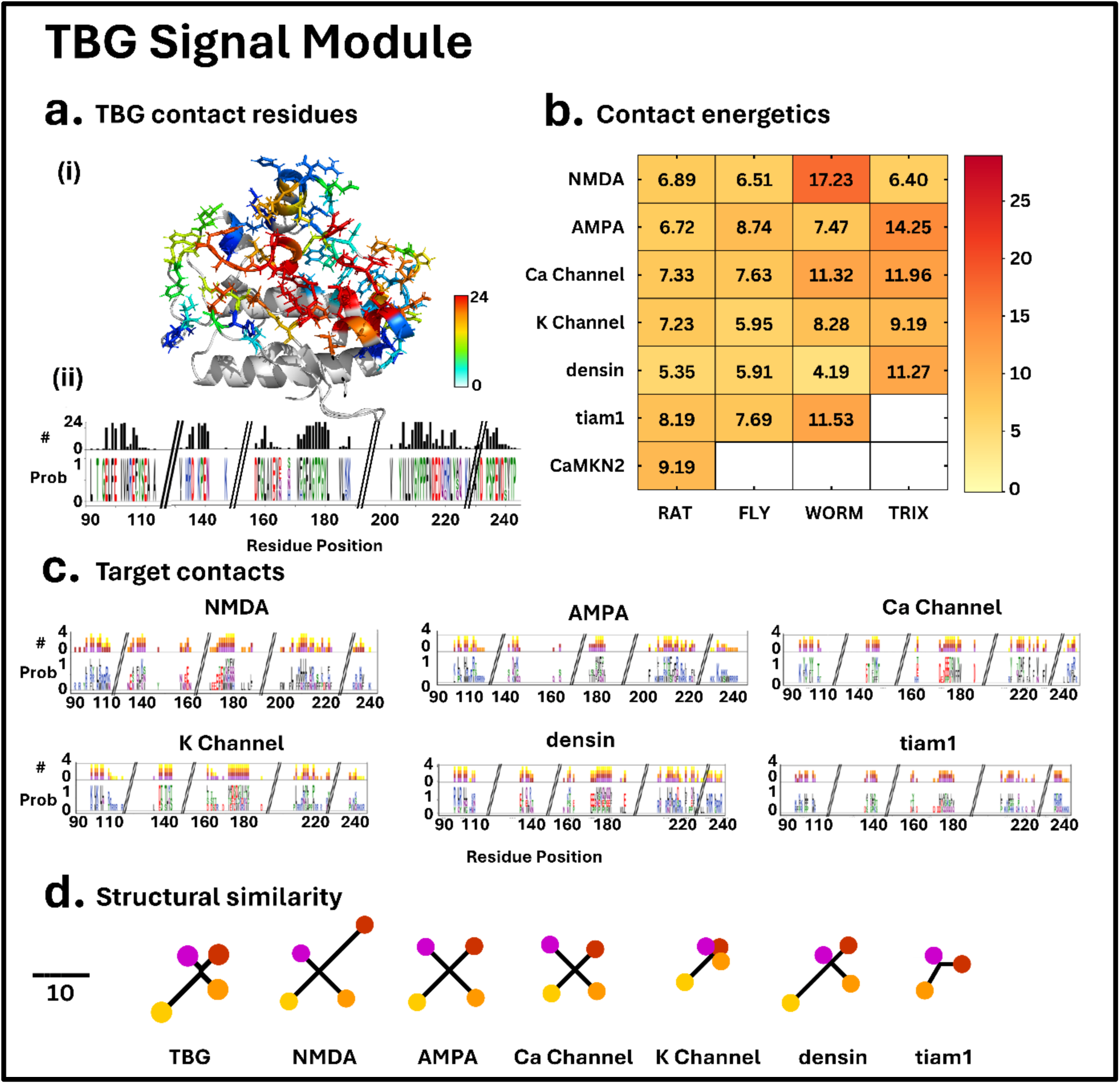
The TBG signal module: **a. Contact residue conservation and target propensity. (i)** Contact density per residue mapped onto structure**. (ii**) HMM logo. Bars (height = number of contacts per residue position) above the HMM logo indicate residue contact distribution. Maximum is 24 (4 species x 6 targets). **b. Contact energetics**. Computed by ePISA. **c. Target contact distribution**. **d**. **Structural similarity.** DALI tree of the structural similarity of the contact segments between organisms for the KD C-lobe (TBG) and its target contact segments. Color code for organisms is as in Fig. 1c.

The CaMKII C-lobe fold that contains the TBG groove has comparable conservation to its targets (**Fig.3d**). Among the ion-channel targets, the K^+^ channel is the most conserved, and the NMDA and AMPA channels are the least conserved. The predicted FLY NMDA is an outlier within the NMDA fold tree. The predicted TRIX representatives have a more divergent fold relative to the other organisms. A tiam-1 TRIX homolog was not found. The FrustratoEvo profiles for the TBG targets are reported in (**Fig. S2d**). No frustrated residue positions that were stabilized by the contact were detected. Thus, the CaMKII contact does not determine the evolution of these targets, primarily ion channels.

### E. The expression profiles support a primordial role for CaMKII in neuropeptide signalling

We downloaded the RefSeq expression profiles of CaMKII and its binding target isoforms for all organisms to relate the biochemistry predicted from the AF complexes to cellular localization (**Fig.4a-d**). In addition to CaMKII, we reported the tissue/cell type distribution profiles of the target isoform with the best match with the CaMKII profile as determined by the Pearson correlation coefficient (*P_cc_*).

**Figure 4:**
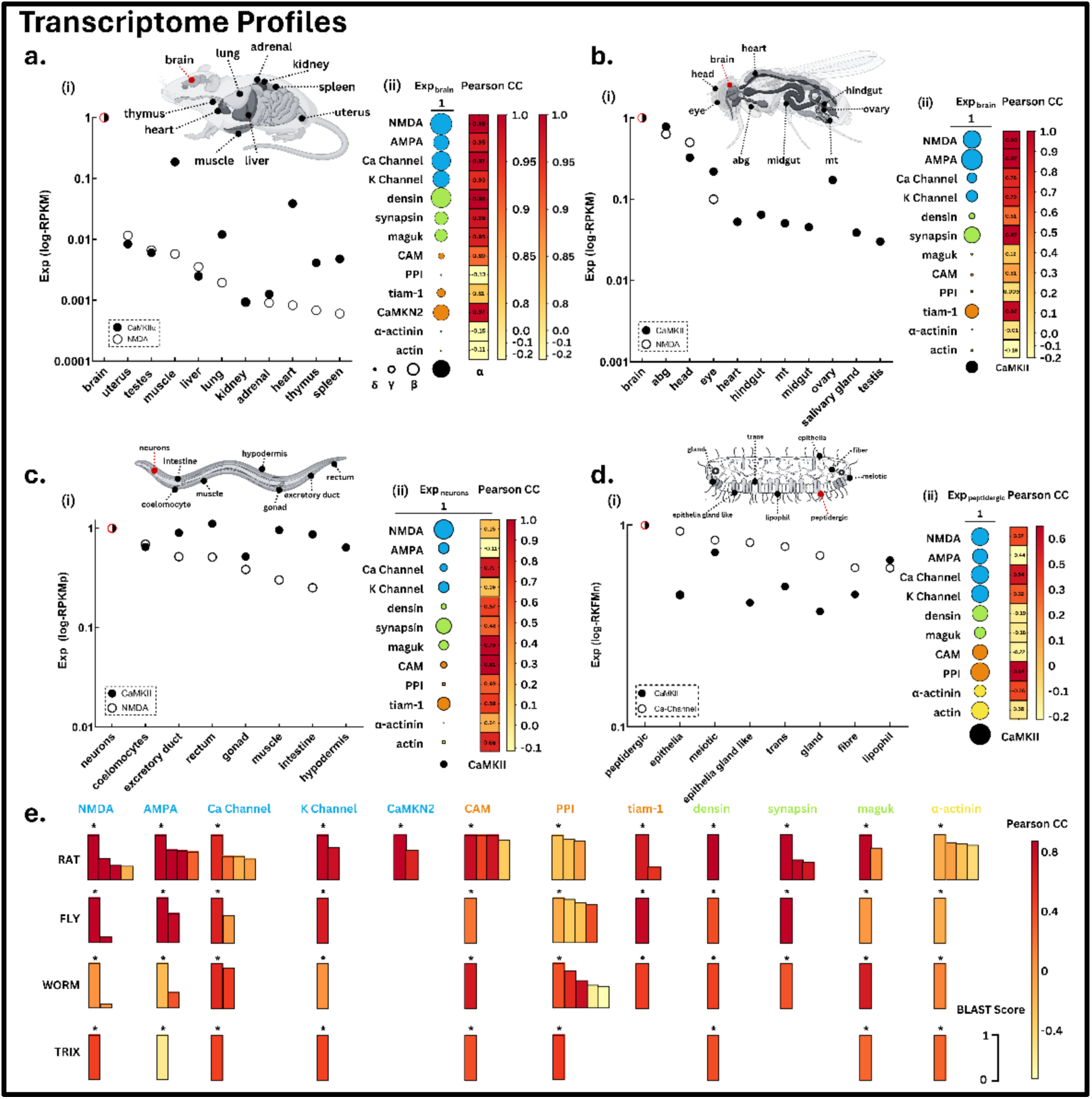
**(a)-(d). The cell/tissue distribution of CaMKII and its target transcripts. (i)** Subpanels show CaMKII RNA-seq expression, as assessed by the RPKM (RAT, FLY), RPKMp (WORM) and RKPMn (TRIX) (**Methods**) across tissue types, normalized by its abundance in neurons (RAT, FLY, WORM) or peptidergic cells (TRIX). The CaMKII distribution is compared with similarly normalized distributions for NMDA (RAT, FLY, WORM) and the L-type Ca^2+^ channel (TRIX). **(ii)** (Left) Subpanels show the abundance of CaMKII (black circle) and its targets (cyan (ion channel), orange (signal), green (synaptic scaffold), yellow (cytoskeletal) across tissue types. The abundance is normalized by division with the sum total of the transcripts per organism (horizontal bars). (Right) Listed, color-coded Pearson correlation coefficients (P_cc_) for the CaMKII / target colocalization. Color key (adjacent bars). **(e)** Gene homolog distributions for all 11 targets within each organism. Homolog counts / organism (RAT (n=35), FLY (n=19), WORM (n=18), TRIX (n=9)). Bar height indicates normalized BLASTp score with top homolog (asterisk). Bar fills indicate Pcc values for the CaMKII / homolog colocalization. Color key (adjacent bar).

In the RAT, the α isoform had the highest neuronal expression ratio, consistent with earlier studies. It, like the NMDA receptor, is expressed almost exclusively in the brain (**Fig.4a**).

In the FLY, the NMDA receptor continued to be the neuronal marker expressed throughout the brain, the abdominal ganglia (abg) and the eye. Its expression in other organs/tissues was <0.001%. There was significant CaMKII expression in the ovaries, consistent with CaMKII ovulation via the octopamine receptor in the FLY oviduct ^58^. CaMKII was expressed in all FLY cell types over a narrower (1 -> 0.01%) range, normalized to its level in the brain, as opposed to the RAT (**Fig.4b**).

In the WORM, NMDA remained the dominant neuronal marker, but its expression was broadly distributed over tissues (1 -> 0.1%). CaMKII expression was even more broadly distributed, with levels in muscle, the intestine, the excretory duct and rectum matching or exceeding that in neurons. In addition, CaMKII expression is most closely correlated with Ca^2+^ channels, rather than NMDA, across tissue types, in contrast to the RAT and FLY (**Fig.4c**).

The lack of synapsin and the Tiam-1 exchange factor reflects the absence of synapses and simpler intracellular circuitry in TRIX. CaMKII expression correlates best with Ca^2+^ channels, amongst the ion channel targets and the PPI phosphatase. CaMKII and Ca^2+^ channels are both most abundant in the peptidergic cell types that mediate neuropeptide secretion. There is less than a 2-fold variation in the expression range over cell types for both CaMKII and Ca^2+^ channels (**Fig.4d**).

Like CaMKII, a number of its binding targets had multiple isoforms (**Fig.4e**). A more complete description of Isoform diversity that includes alternatively spliced variants is documented in **Fig.S3**. There is an increase in both gene and splice variant diversity during the transition from early metazoans to mammals. We finally evaluated the probability of target interactions with CaMKII to evaluate the effect of diversity on the CaMKII interactome (**Fig.S4**).

## Conclusions

The computational modelling of protein-protein interaction networks has emerged as a major research area based on high-throughput genomic and proteomic experiments allied with computational tools. Studies of protein-protein interactomes at atomic resolution required for molecular therapeutics or evolution remain sparse. A notable exception was the work of Mosca et.al.^59^ that used Modeller to model atomic complexes. Here, we have extended their approach and our work on the energy frustration, residue coevolution of the CaMKII hub with the AlphaFold neural network algorithm. Residue coevolution can decipher multiple conformations^60,61^. It is an input for the AlphaFold algorithm^26^, in contrast to Modeller. This study presents the first construction of CaMKII interactomes at the atomic level over four divergent organisms.

### CaMKII architecture

The atomic models, first, addressed the following core issues regarding CaMKII functional architecture - the assembly of the holoenzyme, the occlusion of the TBG by the R regulatory helix and the dissociation of this contact by Ca^2+^.CAM. The binding energy of the CaM-R complex is 2-fold greater than observed for other CaMKII binding proteins, and the survey of the Ca^2+^.CAM / CaMKII complexes suggest that their architecture is conserved across the four organisms. The atomic models of CaMKII complexes with other proteins reveal that energy frustration of the CaMKII R helix and the TBG residues required to maintain their pluripotential binding interfaces is a common feature. Specifically, the R-TBG contact is stabilized by the association when the CLOSED state is matched with dissociated Rf and OPEN C-lobe.

### De novo models

A subset of the AF models evaluated experimental structures of the bound complexes. For example, the Ca^2+^.CAM / CaMKII complex for 2WEL.pdb. Many more models were *de novo* complexes based on biochemical knowledge, but without any structural representative in the four organisms. These included R complexes with PPI, synapsin, α-actinin and Maguk plus the TBG / Ca^2+^ channel complex. The TRIX interactome was based entirely on *de novo* AF models. Successful construction of the complexes validated the biochemical evidence for their existence. In turn, the atomic models extend the mechanistic analysis of CaMKII interactome evolution

### The timing of the Ca^2+^ response

The CaMKII Ca^2+^ response is set by the competition between Ca^2+^.CAM and PPI for R binding sites. Computational models have studied the role of PPI-mediated dephosphorylation in CAM association as it relates to frequency tuning, bistability and cooperativity. This role hinges on the competition between CAM and PPI for a common binding surface^43^. The AF models reveal, for the first time, the atomic details of this surface. We have tracked the relative affinities of these competing targets during CaMKII evolution. The CAM footprint on R*f* includes the inhibitory T305.T306 phosphorylation sites. The PPI footprint does not. Instead, it straddles the inhibitory and primary (T286) phosphorylation sites competent to dephosphorylate T286 in autonomously activated CaMKII. The affinity difference between PPI and Ca^2+^.CAM implies that PPI displacement of Ca^2+^.CAM will depend on Ca^2+^ concentration. As opposed to PPI, which would uniformly distribute in the cytosol, synapsin, α-actinin and Maguk are likely to be localized in cortical membrane microcompartments with substantially greater local concentrations. The energy frustration profiles indicate that maintenance of the R*f* contact influences CAM and PPI evolution, but not the other targets. Energy frustration for the CaMKII-CAM complex was studied previously in the pre-AF era, as part of a larger study of sixty CAM-target complexes with the focus on CAM frustrated residue positions^62,63^. The CAM energy frustration reported here with the FrustratoEVO tool is consistent with these studies.

### The conservation of the TBG and its binding partners

The TBG extends through much of the C-lobe. Its contact residues are well conserved in sequence, as well as the overlap between its target peptides. The target peptides are in the extended conformation and have comparable fold variation. The computed contact energetics are comparable to the R*f* targets except for CAM. The regulation of cytosolic signal transduction, based on the presence of CaMKIIN and tiam1, is most stringent in the RAT and the least in TRIX. The other targets are all membrane channel subunits. The probability that they form complexes with CaMKII will depend on their tissue-specific expression levels and distribution, whether uniform or punctate, in the membrane. If localized as puncta, they would be anticipated to form supramolecular scaffolds as has been proposed for the mammalian CaMKII-NMDA complex^4^. The influence of CaMKII complex formation on the evolution of the target contact sequences could not be detected with energy frustration analysis.

### The TRIX CaMKII interactome for neuropeptide secretion

TRIX has voltage-gated action, sodium action potentials ^64^ and cilia-mediated chemotactic behavior ^65,66^. The overview of the TRIX CaMKII interactome indicates that complexes between CaMKII and its synaptic targets formed before the emergence of synapses. TRIX lacks machinery (synapsin, Tiam-1) for fine-tuned regulation of response timing, and there would be no need for the formation of stable scaffolds for long-term effects. The predicted TRIX CaMKII CLOSED state, and hub architecture are competent to respond directly to Ca^2+^ channel-triggered Ca^2+^ oscillations mediated via Ca^2+^.CAM. In peptidergic cells, the expression of all four channels (NMDA, AMPA, Ca^2+^, K^+^) transcripts is comparable, but the Ca^2+^ channel is most abundant, and its expression correlates most closely with CaMKII. This evidence implies that the dominant TRIX CaMKII function is neuropeptide secretion mediated dominantly by Ca^2+^ channels.

Interestingly, the WORM CaMKII interactome seems to be an intermediate in the transition between neuropeptide and electrical signalling. CaMKII transcripts are broadly distributed across tissue types, consistent with evidence that neuromuscular coordination in the WORM intestine and excretory duct requires CaMKII^67^. The subsequent emergence of mammalian isoforms exploited the subsequent explosion of tissue types to access large variations in expression levels for specialized tasks. Thus, we propose that the CaMKII structure was entrained throughout to sensory signalling, whether via neuropeptides or electrical synapses. The increasing complexity of the body plan and tissue diversity led to the CaMKIIα isoform, singularly dedicated to mammalian cognitive function.

### Limitations of the study

There is increasing evidence from other studies, particularly proteins with switchable folds^68^, that AlphaFold has a limited ability to predict conformational plasticity. Examples of this limitation in this study were predictions of (i) a single Ca^2+^-bound CAM conformation, even though alterations in Ca^2+^-occupancy elicit large changes in the CAM fold ^62^, (ii) a single CAM-bound CaMKII complex, whereas it is likely that there are at least two, weak and strong, binding Ca^2+^.CAM binding states, while apo-CAM does not form a complex ^69,70^, (iii) a single closed ring conformation of the TRIX CaMKII tetramer, whereas both closed and spiral structures have been reported for the closely related cnidarian, *N. vectensis*.

We have restricted our study to binary complexes formed with single CaMKII subunits, given the limitations illustrated by these examples. Predictions of ternary complexes, of possible accessory AD or KD N-lobe binding sites or holoenzyme complexes that form supramolecular scaffolds merit study, once improvements to the AF algorithm to accommodate conformational plasticity in response to covalent modifications, metal cofactors and environmental parameters (e.g. pH) have been established. In the meantime, the insights from the present study into the natural mutations for CaMKII interactome evolution provide an initial platform for therapy of the mutations that underlie CaMKII-mediated disabilities.

## Methods

### Databases

Resource databases for 1D-sequence, 3D-structure, genomes and transcriptomes are listed in **Table 1 (section A)**. Online webservers for sequence and structure analysis are listed in **Table 1 (section B)**, and in-house software/custom code in **Table 1 (section C)**. The metrics for the tissue-specific RNA-seq expression levels for its targets differed slightly. The standard RPKM (reads per kilobase per million mapped reads) normalizes for RNA number and the total number of reads in the measurement ^71^. For WORM expression levels, a prediction tool for 75 distinct tissues ^72^ utilized a training set of RNA-seq samples integrated with protein expression annotated by imaging GFP-labelled cell types (RKPMp). For TRIX expression levels, cell types were identified with a multivariate feature selection algorithm ^73^. Gene expression levels, reported by RNA-FISH and peptide mass spectrometry, were subsequently normalized as the geometric mean for each cell type divided by the median value obtained across all cell types (RKPMn).

### AF model construction

AlphaFold (AF) models were generated on the Biowulf supercomputer (https://hpc.nih.gov/) with AlphaFold2 ^26^. The limitations of AF-models in modelling loop regions, multiple conformational states and allosteric effects pose challenges for AlphaFold2 ^74^ and AlphaFold3 ^75^.

The model construction and analysis are outlined in **Fig. 5**. Five experimental structures were selected for validation of AF predictions of CaMKII monomer domains, homoligomers, and heterodimer target complexes. Models predicted by AlphaFold2 and AlphaFold3 performed similarly in these validation tests. AlphaFold2 was primarily used, as the benefits of AlphaFold3 ^25^ in small molecule and protein nucleic acid complex prediction were not relevant. AlphaFold2 produced 5 models for monomers and 25 models for complexes. The AlphaFold2 (default AF) results are shown (**Table 2**). The model pLDTTs (predicted Local Distance Difference Test profiles) have been deposited (see Data Availability Statement).

**Figure 5:**
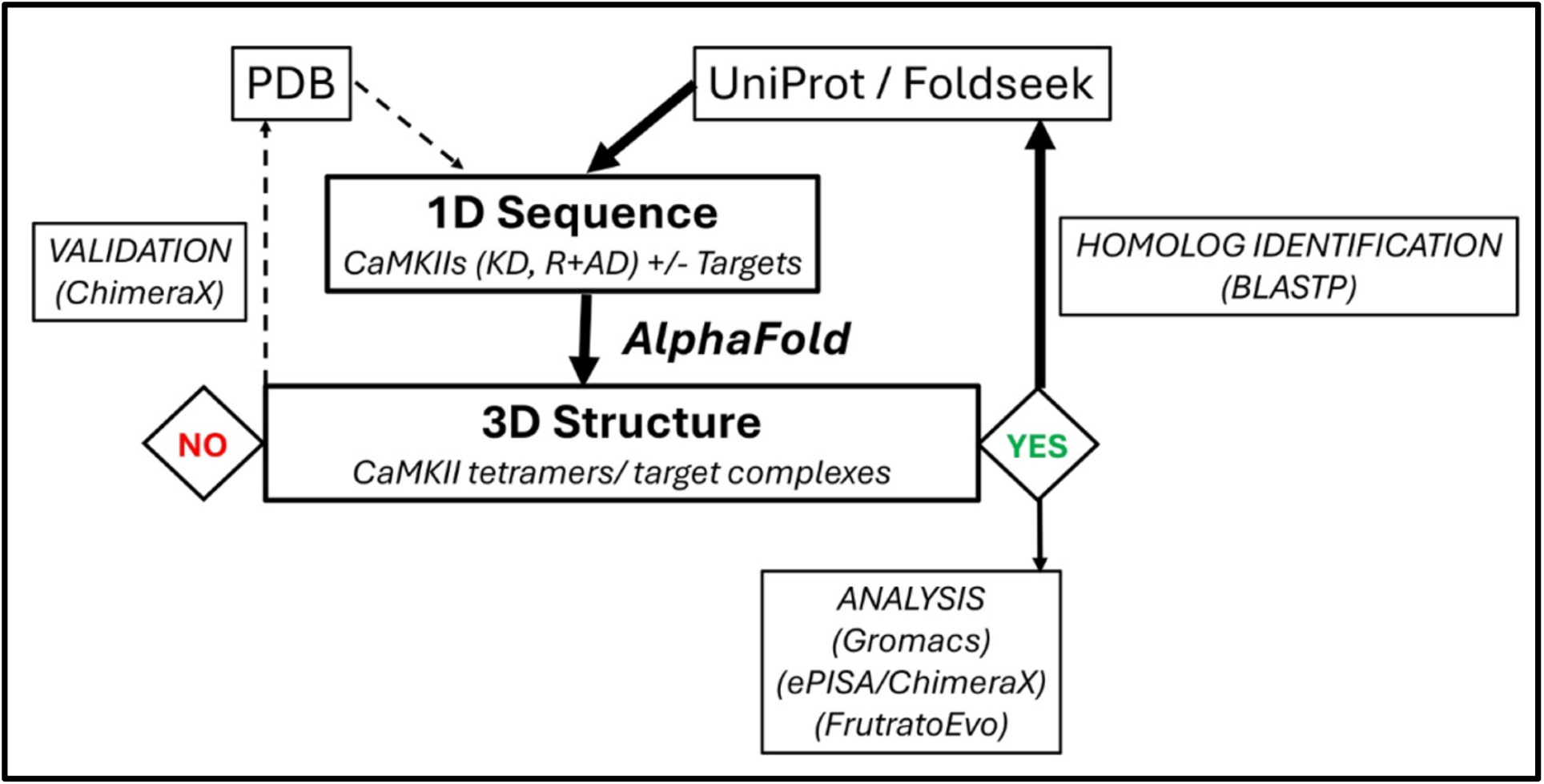
Structural Analysis. Schematic of the AF model generation and analysis pipeline. The top homologs were identified, typically from multiple homologs, by the BLASTP match score / E-value, length (number of residues), and topology of the target sequence in the 3D Foldseek AF-model associated with the target subunit/domain. Once a model was generated, the target sequence was used to identify homologs in the lower organism most closely related in the phylogenetic tree of life. The process was repeated until homologs for all 4 species were identified (thick arrows). Table 2 lists the validation results. The AF models were energy minimized before analysis with short molecular dynamics (MD) simulations (Gromacs). ChimeraX, ePISA and FrustratoEvo were used to study the energetics of the homodimer and heterodimer contacts. Fold evolution was determined with DALI, and sequence conservation with HMMER3 (see text).

CaMKII Models: Sequences for AF model construction of AD tetramers, CLOSED and OPEN state KDs were selected by homology to the RAT CaMKIIα sequence. Analysis of the CaMKII linkers, restricted to 1D sequence, proceeded iteratively with RAT linker variants identified relative to the reported human linkers, then used to identify FLY linker orthologs. The process was iterated (RAT -> FLY, FLY -> WORM, WORM -> TRIX). The BLASTp match scores were compared against a decoy match score distribution as well as the maximum possible score obtained by self-match (**Fig. S1**).

Homolog identification: Target RAT sequences were selected first based on BLASTp matches in UniProt to sequences identified from crystal structures of CaMKII complexes or biochemical assays (see **Introduction**). These target sequences were used together with the RAT CaMKII KD-R or R+AD sequences as inputs for AF model construction. Inspection of the contact maps revealed that the top model typically had a very similar confidence profile to the next five top-ranked complexes. The RAT target sequences were then used for identification of the corresponding target orthologs in the FLY genome and so on, with iterative repetition similar to the protocol outlined above for CaMKII linker segments. These sequences were then used with the R or R+AD sequences of the corresponding organism for AF model construction.

In addition, the target proteins represented in the AF complexes were used as probes for paralog identification with a UniProt BLASTp search within their genome. Identified gene and alternatively spliced variant isoforms were distinguished by an MSA to detect spliced segments. Finally, the target contacts predicted by the AF complexes were used to probe the MSA to identify possible CaMKII targets. In all cases, the matches followed the protocol established from analysis of the CaMKII linkers

Two criteria were used to validate and characterize target contacts. First, the predicted contacts were compared against homologous contacts in experimental 3D-structures. In the absence of structural data, they were checked for agreement with contacts inferred from biochemical protection or crosslinking data, if available. The targets for the mammalian RAT complexes were selected first because the predicted models could be evaluated based on either the first or second criterion. The AF models for the other organisms were built from homologous sequences found from the iterative search protocol noted above. The RAT CaMKII interactome was built with 11 AF-model complexes. The interactome size decreased in the lower organisms due to the loss of interactome components from their genomes. HMM logos mapped position-dependent sequence conservation between CaMKII from different species.

### AF-Model Analysis

The models were energy-minimized in GROMACS 2021.4 with the CHARMM36 forcefield. These simulations and subsequent analysis of the energy-minimized models were done on a compute node (Dell Power Edge R730, 2 x Intel Xeon E5-2678 v3 (12 Cores), 224 GB RAM (16GBx14), 600 GB RAID+1 TB RAID + NVIDIA Tesla K40C (2880 Cores)) in the in-house BIRL computer cluster.

ChimeraX was used for alignment of structural models and identification of residue-residue contacts (< 5 angstrom contact distance). ePISA was used to determine residue contact energies. The match of the contact residues identified by ChimeraX and ePISA was >90%.

The 3D-fold evolution of CaMKII and its targets was characterized with DALI ^76^. The DALI server uses intramolecular distance matrices to compare 3D-structures ^77,78^. It can detect remote homologs, assisted with hidden Markov models ^79^, in experimental and AlphaFold predicted protein databases ^80^. The structural segments must be longer than 30 residues for DALI fold comparison. This criterion was met for short target segments by expansion of the N and C-termini of the contact segment.

The FrustratoEvo server was built on the Frustratometer algorithm ^55,81^ that we described previously ^82,83^. FrustratoEvo extends the central Frustratometer concept of local frustration as assessed by a decoy distribution of *in silico* mutants to rank the energy stabilization of residue contacts is extended by FrustratoEvo to maps the evolution of the local frustration at the single residue level in protein families.

## Abbreviations

*Ca^2+.^CAM*: calcium calmodulin
CaMKII: Ca^2+.^CAM-dependent protein kinase II
AD: association domain
KD: kinase domain
TBG: target binding groove
R: regulatory helix
AF: AlphaFold

## Author Contributions

**S.T.K.** mined the seqRNA expression databases, performed energetic and contact analysis of the atomic models, and prepared the manuscript figures. **M.H.** wrote custom scripts to perform *in-silico* mutagenesis, assess binding energetics and frustration scores, and contributed to the analyses of the AF atomic models and preparation of the manuscript. **T.S.R.** funded and co-designed the project. **S.K.** co-designed the project, constructed all models, wrote the manuscript, and supervised S.T.K. and M.H.

## Acknowledgements

This research was supported [in part] by the Intramural Research Program of the National Institutes of Health (NIH), National Institute of Neurological Disorders and Stroke (NINDS). The contributions of the NIH author(s) were made as part of their official duties as NIH federal employees, are in compliance with agency policy requirements, and are considered Works of the United States Government. However, the findings and conclusions presented in this paper are those of the author(s) and do not necessarily reflect the views of the NIH or the U.S. Department of Health and Human Services. The AlphaFold models were constructed on the NIH HPC Biowulf computational cluster (http://hpc.nih.gov) and energy-minimized at BIRL, LUMS (https://birl.lums.edu.pk). We are grateful to the LUMS Lifesciences department, and particularly. Professors Muhammed Tariq and Safee Ullah Chaudhary (S.U.C) for support for S.T.K. and M.H., respectively. M.H. was supported, in addition, by the Higher Education Commission (HEC) of Pakistan grant 20-3629/NRPU/14/585 (to S.U.C). We thank Drs. Steven Vogel and Howard Schulman for comments on the manuscript. Dedicated to the late Dr. Thomas S. Reese.

## Data Availability Statement

All AF models have been deposited in ModelArchive ( https://modelarchive.org/). The ChimeraX files of the residue contacts, ePISA files of the contact energetics, and the MSAs for gene and splice variant homolog identification have been deposited in Figshare (https://figshare.com/).

