## Supplementary Material for "An AlphaFold guided model for the evolution of the CaMKII interactome"

### Supplementary Information

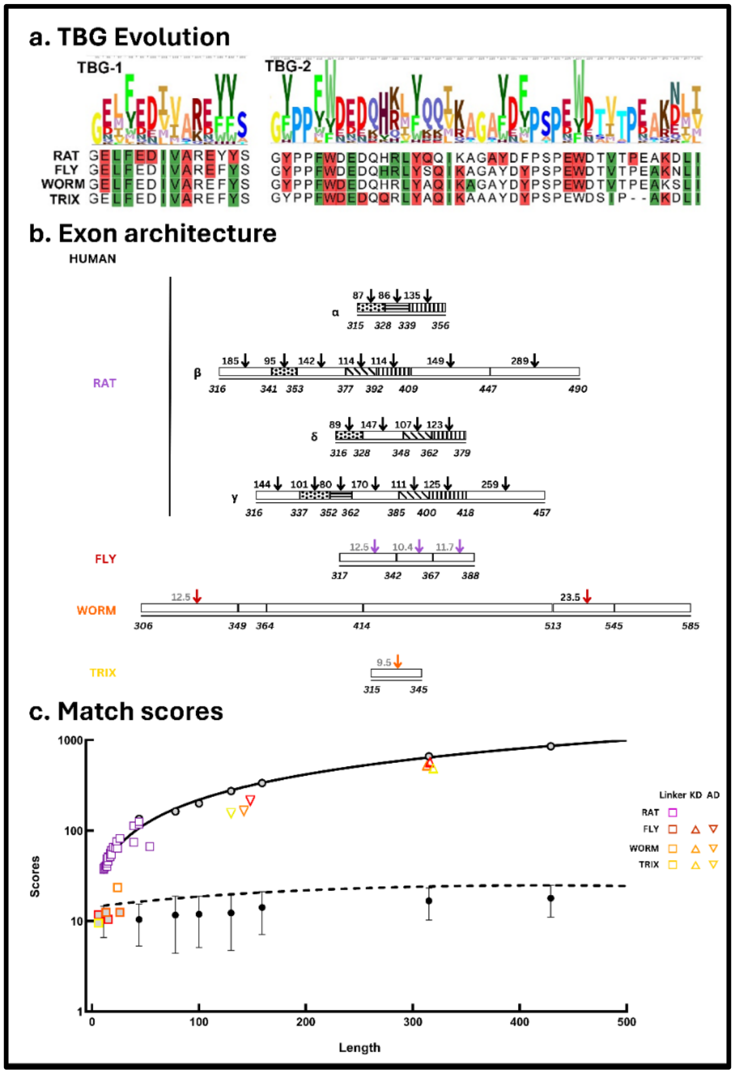


***Supplementary Figure S1: a. TBG energy frustration is not correlated with sequence conservation.*** *(Top) The HMM logo of KD C-lobe residue conservation in 89 species (from [1]). (Bottom) The FrustratoEvo alignment for the 4 organisms showing the maximal (red) and minimal (green) frustrated residue positions.* ***b. Exon architecture of linker domains.*** *Colors indicate organisms, as in Fig. 1c. Homologs for human linker exons [2] were mapped (arrows) to the RAT isoforms. The RAT exons were then used to search for homologous FLY CaMKII linker exons, and so on. Arrows indicate significant match scores (numerical labels). Exon lengths reflect their sequence lengths. Common motifs (e.g. striped) indicate homologous exons across species.* ***c. Match scores.*** *The maximum values for the BLASTP matches were obtained by self-match of the query exon sequences (open symbols) with best-fit (solid line). Randomly-generated 100-sequence decoy distributions at various target lengths filtered out random matches. Most match scores fell in the range between the maximum match score and the* $+3\sigma$ *threshold (best-fir (dashed line) for the decoy distribution. Filtered out short exon matches (light fills). Match scores for the KD and AD (Fig. 1a) are shown for comparison.*

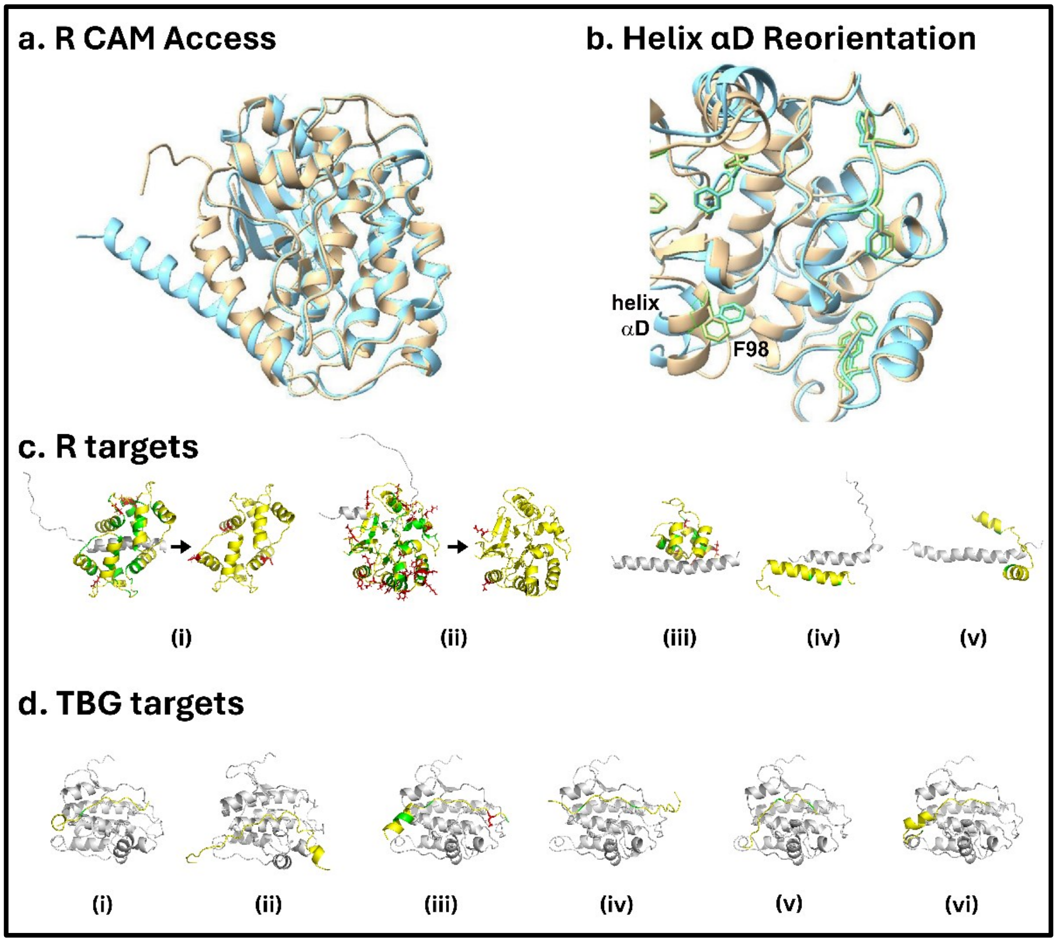


***Supplementary Figure S2: a. R undocked by CAM.*** *The alignment of the CLOSED KD C-lobe state with docked R (gold) and the OPEN state (cyan) created by CAM association reveals the displacement of the R helix from the C-lobe core. The CAM is not shown.* ***b****.* ***Reorientation of helix $\alpha$D by bound substrate****. The alignment of the CLOSED KD C-lobe and the OPEN state formed when the GluN2B subunit is bound. The rotation of the F98 sidechain (asterisk) is a diagnostic for the rotation of helix $\alpha$D[3], replicated in these AF models of the RAT complexes. The docked R-helix (CLOSED state) and the bound GluN2B (OPEN state) are not shown.* ***c, d Target FrustratoEvo profiles.*** *Residue positions with maximal (red) and minimal (green) frustration are mapped onto the RAT target complexes of R targets.* ***c.*** *R targets. i. CAM, ii. PPI, iii. $\alpha$-actinin , iv. Synapsin-1, v. Maguk.* ***d.*** *TBG targets. i. AMPA, ii. NMDA, Iii. Ca2+ channel, iv., K channel. v., Densin. vi. Tiam-1.*

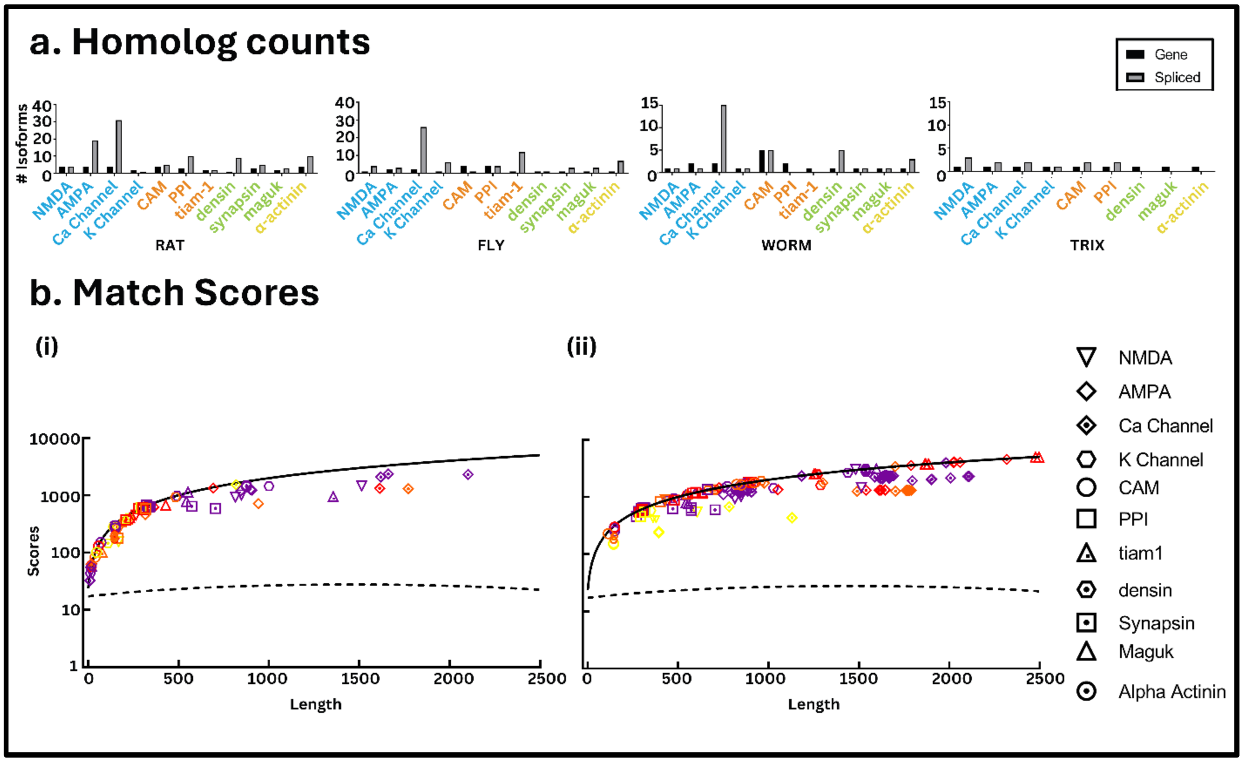


***Supplementary Figure S3: a,b. Homolog Diversity of CaMKII targets. a. Homolog counts.*** *Counts are reported separately for gene homologs (RAT (n=33), FLY (n=19), WORM (n=18), TRIX (n=9)) and alternatively spliced variants (RAT (n=99), FLY (n=70), WORM (n=33), TRIX (n=12))*. ***b. Match scores.*** *For each organism, its genome was probed with the target protein used in the AF complexes for identification of gene homologs and alternatively spliced variants (see* ***Methods****).* ***(i) Gene homologs. (ii) Alternatively spliced variants.*** *The lines represent fits to the maximum score and decoy distributions as in Fig. S1c.*

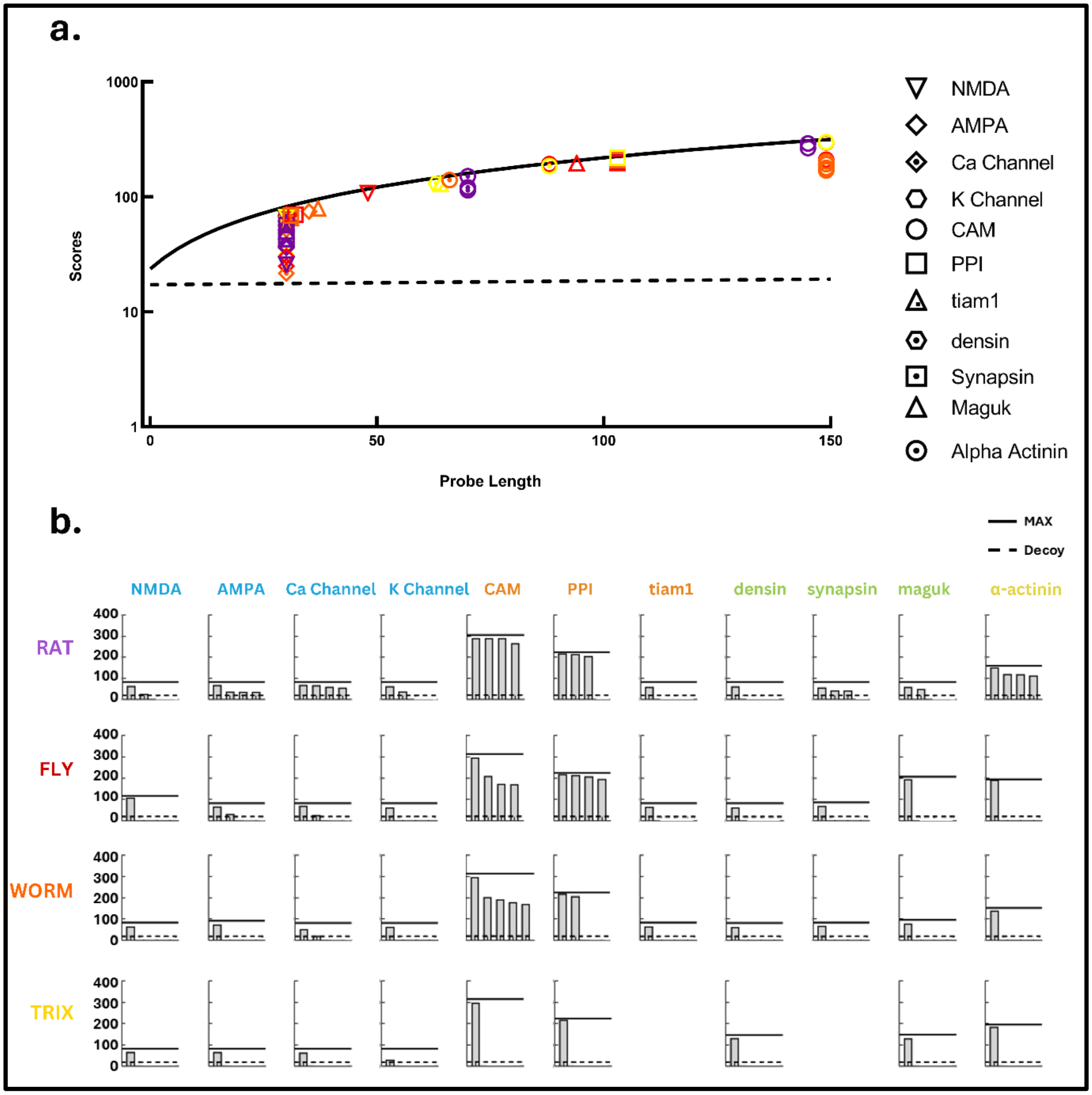


***Supplementary Figure S4: a. Match scores for target contact segments.*** *Distinct gene homologs are denoted by species and target protein as in Fig. 1b.* ***b. Possible CaMKII interactors.*** *The sequences of the target structures in the AF models were used as probes to rank possible CaMKII targets from the identified gene homologs. The solid and dashed lines represent best-fits to the maximum score and decoy distributions as in Fig. S1c. Possible CaMKII targets > decoy score (m+3 σ) were RAT (n=30), FLY (n=19), WORM (n=17), TRIX (n=9).*
